# Letrozole delays acquisition of water maze task in female BALB/c mice: possible involvement of anxiety

**DOI:** 10.1101/2022.11.01.514755

**Authors:** Jacek Mamczarz, Malcolm Lane, Istvan Merchenthaler

## Abstract

Letrozole, an aromatase inhibitor (AI), is used as an adjuvant therapy in estrogen receptor-positive (ER+) breast cancer patients. Similar to other AIs, it induces many side effects, including impaired cognition. Despite its negative effect in humans, results from animal models are inconsistent and suggest that letrozole can either impair or improve cognition. Here we studied effects of letrozole on cognitive behavior of adult female BALB/c mice, a relevant animal model for breast cancer studies. Mice were continuously treated with once-a-day subcutaneous (s.c.) injections of letrozole (0.1 or 0.3 mg/kg/day) or vehicle and subjected to behavioral testing starting on day 21 after treatment initiation. During the treatments, vaginal smears were taken from the mice to evaluate estrous cyclicity. Both doses of letrozole suspended cyclicity and the smears showed that the mice were in constant metestrus. Exposure to letrozole did not significantly affect response to novelty measured as a locomotor activity in open field. However, repeated testing in open field (4 days × 15 min) revealed that letrozole 0.3 mg/kg facilitated locomotor habituation (a form of non-associative learning), significantly reducing locomotor activity on 3rd and 4th day of testing. These findings suggest that certain doses of letrozole may have positive effects on cognitive behavior. Training to find a hidden platform in the Morris water maze (15 days x 4 trials), however, indicated that letrozole 0.1 mg/kg-treated mice had significant learning impairment, as, throughout the training, they swam longer times than vehicle-treated mice to reach the hidden platform. Similarly, in a probe test performed 72 h after the last day of the training, letrozole 0.1 mg/kg-treated mice did not show preference for the training platform zones. These results indicate that cognitive impairments reported by women treated with letrozole can be captured in BALB/c mice treated with clinically relevant doses of the drug. Interestingly, most of the letrozole 0.1 mg/kg-treated mice were able to learn the new platform position in reversal training and performed similar to control mice in a reversal probe test. Results of the reversal test suggest that letrozole did not completely disrupt spatial navigation but rather delayed acquisition of spatial information. The current study shows that letrozole dose dependently modulates behavioral response and that its effects are task dependent.

## 1. Introduction

Breast cancer is the most common cancer in women (Ferlay et al., 2015), with nearly 70% of patients having estrogen receptor-positive (<u>ER+</u>) cancers. Women suffering from ER+ breast cancers are commonly treated with aromatase inhibitors, which, by inhibiting the activity of the enzyme aromatase (Burstein et al., 2014) encoded by the *CYP19* gene, prevent the synthesis of 17β-estradiol (E2) from testosterone (T). Aromatase inhibitors, such as letrozole, are usually prescribed as daily oral treatment over 3 to 5 years (American Cancer Society, 2019). However, these drugs have long been associated with CNS-related side effects that impair quality of life, including hot flushes (Rand et al., 2011), cognitive deficits (Collins et al., 2009; Jenkins et al., 2004; Schmidt et al., 2000), insomnia (Desai et al., 2013), and depression (Ganz et al., 2016). Importantly, these side effects can become so severe that a significant proportion (~ 15 %) of women discontinues this potentially life-saving treatment (Chirgwin et al., 2016). Thus, there is an urgent need to introduce novel therapies that can mitigate these CNS-related effects and, thereby, improve the quality of life of breast cancer patients treated with aromatase inhibitors to increase treatment compliance.

The CNS-related side effects of aromatase inhibitors are thought to be due to lack of E2 produced *de novo* in different brain regions, including those associated with learning and memory, regulation of sleep, depression, and anxiety (e.g., hippocampus, cerebral cortex, amygdala, and raphe nuclei) (Roselli et al., 1985; Stoffel-Wagner et al., 1999). Indeed, the *de novo* synthesis of E2 in the hippocampus induced by aromatase results in higher E2 concentrations (8.4 nM) in the hippocampus than in the peripheral blood (0.014 nM) (Hojo et al., 2009).

The hippocampus has a central role for memory formation and consolidation in the mammalian brain (Mukai et al., 2010), and hippocampal functions are highly supported by E2. Dendritic spine and synaptic densities are regulated by the estrus cycle (being the highest in proestrus) (Woolley and McEwen, 1992), E2 deficiency (induced by ovariectomy [OVX]), and OVX/E2 replacement (Kaidah et al., 2016; Li et al., 2004; Tuscher et al., 2016a; Woolley et al., 1990; Woolley and McEwen, 1992; Woolley et al., 1996; Xing et al., 2018). Estrogen increases: (i) synaptic protein levels (such as spinophilin, a marker of dendritic spines), (ii) phosphorylation of Cofilin (p-Cofilin stabilizes the spine cytoskeleton) (Kretz et al., 2004; Vierk et al., 2012; Zhou et al., 2004), and (iii) and downregulates steroid receptor coactivator-1 (SRC-1) expression (Bian et al., 2014; Liu et al., 2015) and the expression of the excitatory synapse marker PSD95 (Liu et al., 2015) each involved in the regulation of hippocampal synaptic plasticity and spatial memory (Bian et al., 2014; Liu et al., 2015). Synapse density and dendritic spine density greatly contribute to synaptic plasticity and to learning and memory behaviors (Dominguez-Iturza et al., 2016; Meng et al., 2002). Hippocampus-dependent spatial reference memory fluctuates with the estrous cycle (Frick and Berger-Sweeney, 2001) and is improved by E2 on OVX animals (Kaidah et al., 2016; Li et al., 2004). Earlier studies demonstrated that intrahippocampal administration of letrozole impaired spatial working memory in intact and OVX rats (Marbouti et al., 2020), and non-spatial memory in OVX mice (Tuscher et al., 2016b). Similarly, inhibition of hippocampal aromatization decreases spatial learning and memory in male zebra finches (Bailey and Saldanha, 2015; Bailey et al., 2017). The detrimental effect of letrozole on cognition appears to be a result of suppressed synaptic plasticity, because inhibition of E2 synthesis by letrozole applied to primary hippocampal cultures or microinjected into the hippocampus of OVX mice reduces the number of dendritic spine synapses (Fester et al., 2012; Hojo et al., 2004; Kretz et al., 2004; Prange-Kiel et al., 2006).

In contrast to intrahippocampal injections, systemic administration of letrozole that is clinically relevant shows inconsistent effect on spatial learning and memory in mice and rats, ranging from impairment (Zameer and Vohora, 2017, Liu et al., 2019) or no effect (Meng et al., 2011;(Vierk et al., 2015) in mice, to improvement in rats (Alejandre-Gomez et al., 2007, Aydin et al., 2008). In addition to differences in the response to letrozole in rats and mice, there are differences among mouse strains (Cestari et al., 1999; Ciamei et al., 2000; Lipartiti et al., 1993) that can affect the effects of letrozole on cognitive performance. Inbred and outbred strains differ from one another in genetic makeup, in neurochemistry (Ingram and Corfman, 1980), synaptic plasticity, and learning and memory (Nguyen et al., 2000). Several studies have demonstrated strain-dependent effects of GABA-receptor agonists and antagonists in a number of behavioral tests (Castellano et al., 1993; Jacobson and Cryan, 2005; Puglisi-Allegra et al., 1981; Simler et al., 1982). Therefore, comparison of different mouse strains provides valuable information about the importance of genetic background in behavioral and pharmacodynamic phenotypes (Jacobson and Cryan, 2005). To date, C57Bl mice were the most often used strain in studies evaluating effect of letrozle on spatial learning (Meng et al., 2011; Liu et al., 2019; Vierk et al., 2015). Since our goal was to evaluate the effect of prolonged exposure to letrozole on cognitive functions in an animal model used in breast cancer research (Orlandella et al., 2021), the present study used female BALB/c mice. The BALB/c mice are significantly different form C57BL mice in their emotionality (Brinks et al., 2007) and neurochemistry (Bach et al., 2011), therefore their evaluation may provide additional information about effects of letrozole on cognitive behavior.

In this study, intact female BALB/c mice were exposed to a chronic treatment of different doses of letrozole. Then, locomotor activity and locomotor habituation, a form of non-associative learning, were assessed in open fields, while spatial learning and memory were assessed in Y-maze and the Morris water maze (MWM). Data presented here support the hypothesis that similarly to humans, BALB/c mice exposed to letrozole develop learning and memory deficits depending on dose ranges and the specific tasks. Based on the results of this study, the BALB/c mice emerge as a preclinical model for future investigation of countermeasure against side effects of the letrozole therapy.

## 2. Materials and Methods

### 2.1. Animals

Female BALB/c (Charles River Laboratories) mice arrived in the animal facility as 2–3-month-old, weighing 20 ± 2 g). Mice were housed 4-5 per cage with food and water available *ad libitum*, in 12-h light dark cycle (7:00 am–7:00 pm). Mice were subjected to the experimental procedures after one week of acclimation. All experimental procedures were approved by the University of Maryland Baltimore Animal Care and Use Committee and were conducted in accordance with the National Institutes of Health Guide for the Care and Use of Laboratory Animals

### 2.2. Drugs and administration

Letrozole (Sigma) was dissolved in 0.1% DMSO in physiological saline and was administered subcutaneously (s.c.) in a volume of 100 μl. For the drug administration, mice were transported each day to the procedure room.

### 2.3. Vaginal smears

Estrous cyclicity was evaluated on a daily base starting from the second day following the letrozole administration. Evaluation was conducted until mice stopped cycling, and thereafter for an additional week to confirm their estrous status.

### 2.4. Behavioral assessment

#### 2.4.1. Experimental design/timeline

Prior to starting behavioral testing, mice were treated once a day with letrozole or vehicle for 3 weeks to mimic the long-term clinical use of this drug by breast cancer patients. Treatment continued during the period of behavioral testing, with letrozole or vehicle being administered afternoon after completing testing.

Twenty-two days after initiation of treatment, locomotor habituation was tested in open fields (OF) for 4 consecutive days. Then, on the 26^th^ day, spontaneous alternation was tested in the Y-maze. On the 29^th^ day, water maze testing was started. Reference memory training was conducted for three consecutive weeks and was followed by a probe test 72 h later. Then, mice were subjected to reversal learning on 5 consecutive days followed by a probe test 72 h later. Twenty-four hours after the last probe test, mice received additional reference memory training in a new room (two blocks of 4 days interspaced by 72-h interval.

#### 2.4.2. Locomotor habituation in OFs

A day before OF testing, mice were brought to the experimental room for 2 h of habituation, after which time they were exposed for approximately 10 min to a cage the same as a housing cage but with bare floor. Immediately before testing in the OF, all mice tested at the same time, were put singly to the cages with bare floor for around 5 min, and then were caried in these cages to the testing apparatus. Up to 12 mice were tested simultaneously in 12 OFs (60 × 60 × 60 cm), made of black Starboard plastic. Light intensity in the center of each open field was 20 Lux. Conair sound machines were set to generate “white noise” on one side and “running stream” sound on the opposite side of the apparatus. Mice were tested for 15 min in each of 4 consecutive days.

#### 2.4.3. Spontaneous alternation in Y-maze

A day after completing OF testing mice were allowed to explore a Y-maze (arm: 9 cm × 9 cm 37 cm) for 10 min under dim light condition (30-35 Lux). The custom build maze had gray-painted wooden walls and was placed on black Starboard plastic floor. The maze was cleaned by MB-10 and dried by fan between tested mice.

#### 2.4.4. Morris Water Maze (MWM)

##### General water maze training procedure

Our pilot experiment showed that female BALB/c mice, when tested in 24°C, a standard temperature used for male BALB/c mice (Chapillon and Debouzie, 2000; Huang et al., 2012b; Philpot et al., 2016), were rapidly losing body temperature, were shivering, and were not able to effectively swim in the subsequent trials. Therefore, we applied a procedure by gradually decreasing water temperature over the course of the training that we successfully have used for guinea pigs (Mamczarz et al., 2011). Such approach reduces floating behavior of BALB/c mice that is induced by cold water. BALB/c mice, in contrast to C57Bl mice, have been reported to float less at water temperature of 30°C than at 20°C during forced swim test (Bachli et al., 2008).

In the current study the room temperature was kept constant at 24.5 ± 0.5°C, while the water temperature was gradually decreased from 28 ± 0.25°C to 21 ± 0.25°C, to provide comfort of swimming as well as motivation for escape. During the 1^st^ week, the water temperature was dropped from 28°C to 25.5 ± 0.25°C on the 5th day of training; during the 2^nd^ week from 24.5 ± 0.25°C to 23.0 ± 0.25°C on the 10^th^ day of training. The 3rd week of the training was started at 22 ± 0.25°C, but we noticed that some good learners started losing interest in a direct escape to the platform, therefore on the 12th day temperature was further decreased to 21 ± 0.25°C and was kept at this value to the last day of reference memory training. Light intensity was set to around 70 lux in the middle of the maze and was brighter outside the maze (70-150 Lux) as recommended for male BALB/c mice (Chapillon and Debouzie, 2000; Huang et al., 2012b).

Training was performed in black plastic tank (120 cm in diameter). A round black platform, 9 cm in diameter, was positioned in the middle of one of the quadrants and was submerged 1 cm under black opaque water. Water was made opaque by black tempera paint. On each day of training, mice were subjected to four 90-s trials with 5-min inter-trial intervals (ITI). On the first training day, prior to the first trial, each mouse was placed onto the submerged platform for 15 s, after which they were placed into the drying cage for approximately 5 min prior to starting the first training trial. During the training, mice were placed on the water facing the wall and were allowed to navigate for 90 s to find the platform. After finding the platform, mice were allowed to stay on it for 15 s. If they did not find the platform, initially they were gently navigated by the experimenter, who allowed them to follow a hand in a bright glove. Later, when mice showed preference for the platform, the experimenter pointed the platform by a bright glove and waited until mice climbed on it by themselves. Immediately after being removed from the water, mice were dried using paper towels and placed in warm (approximately 30°C) drying boxes (32 cm × 29 cm × 29 cm) lined with cotton towel for the duration of the ITI.

##### Reference memory learning

Water maze training started on the 29^th^ day of letrozole or vehicle treatment. Mice received 15 days of training in 5-day blocks interspaced by 72-h breaks. Each of four daily trials started from one of four cardinal locations (N, S, E, or W). The water temperature was maintained as described above.

##### Reference memory probe test

The training was followed by a 60-s probe test performed 72 h after the last training trial. The water temperature was kept at 22.0 ± 0.25°C. The temperature was 1.0 °C higher than the temperature kept on the last training day to mitigate stress of cold-water immersion after the three-day break.

##### Reversal learning

Reversal learning started 24 h after the probe test and was continued for 5 consecutive days. The platform was moved to the opposite quadrant, and mice received 5 days of training, 4 trials a day, with 5-min ITI. The starting position was alternated between the two positions equidistant to the previous and the current platform location. The water temperature was kept at 21 °C for the entire training.

##### Reversal probe test

Ninety-second reversal memory probe test was performed 72 h after the last reversal training trial. The water temperature was kept at 22.0 ± 0.25°C

##### Reference memory learning in a new room

The new training started 24 h after completing the reversal learning. Mice received 8 days of training in two 4-day blocks interspaced by 72-h breaks. The training procedure was similar to the one described above in *General water maze training procedure* with except for the water temperature. Water temperature was set to 23 ± 0.25°C on the first day, and then, on the second day, was decreased to 22 ± 0.25°C and was kept on similar level to the end of the training.

#### 2.4.5. Post-MWM open field testing

A few days after completing water maze testing, mice were subjected to testing in large open field (120 cm × 120 cm) for 1 h under dim light condition (20 Lux).

### 2.5. Data analysis and statistics

All data were recorded using the Any-Maze software (Stoelting Co, Wood Dale, IL) and were extracted from the program for statistical analysis. Statistical analysis was conducted using the SigmaPlot 12.0 software. Animal behavior was compared between experimental groups, as well as within experimental group to find within-subject changes in locomotor activity among testing days, or to evaluate within-subject preference to a target training zone versus non-target zones in water maze. Learning curves were analyzed using two-way repeated measure ANOVA. Then, if main effects or interactions were significant, for each testing day, one way ANOVA followed by Tukey *post-hoc* test was conducted. For the probe test, preference to the training target water maze zone was evaluated on three levels of precision: quadrant, annulus-40 (the 40-cm diameter area around the center of the quadrant (Markowska et al., 1993)), and crossings of the target platform area. Among groups comparison was conducted using one-way ANOVA, followed by Tukey *post-hoc* test. Within-subject preference to water maze zones was analyzed using one-way repeated measure ANOVA (RM-ANOVA) followed by Dunnet’s *post-hoc* test, in the following way: for quadrants, time and distance swam in a target quadrant was compared with time and distance in remaining 3 quadrants; for annulus-40, time and distance in the target annulus-40 was compared to time and distance in annuli-40 virtually positioned in the center of non-training quadrants; crossings of the target platform was compared to crossing of areas equal to the platform area that were virtually positioned in the center of non-training quadrants. A schematic representation of the water maze areas is presented in Figure 1. If normality or equality of variance was violated, an equivalent non-parametric test was conducted. Specifically, one-way ANOVA was replaced by Kruskal-Wallis One Way Analysis of Variance on Ranks (ANOVA on Ranks), and one-way RM-ANOVA was replaced by Friedman Repeated Measures Analysis of Variance on Ranks (RM-ANOVA on Ranks). Tukey *post-hoc* test was replaced by Dunn’s post-hoc test for the non-parametric comparison. Dunnet’s *post-hoc* test was used after within-subject comparison analyzed by RM-ANOVA or RM-ANOVA on Ranks. Significant difference detected with *post-hoc* tests are shown as p<0.05, because this level of significance is only presented by SigmaPlot 12.0 for the non-parametric tests.

**Figure 1.**
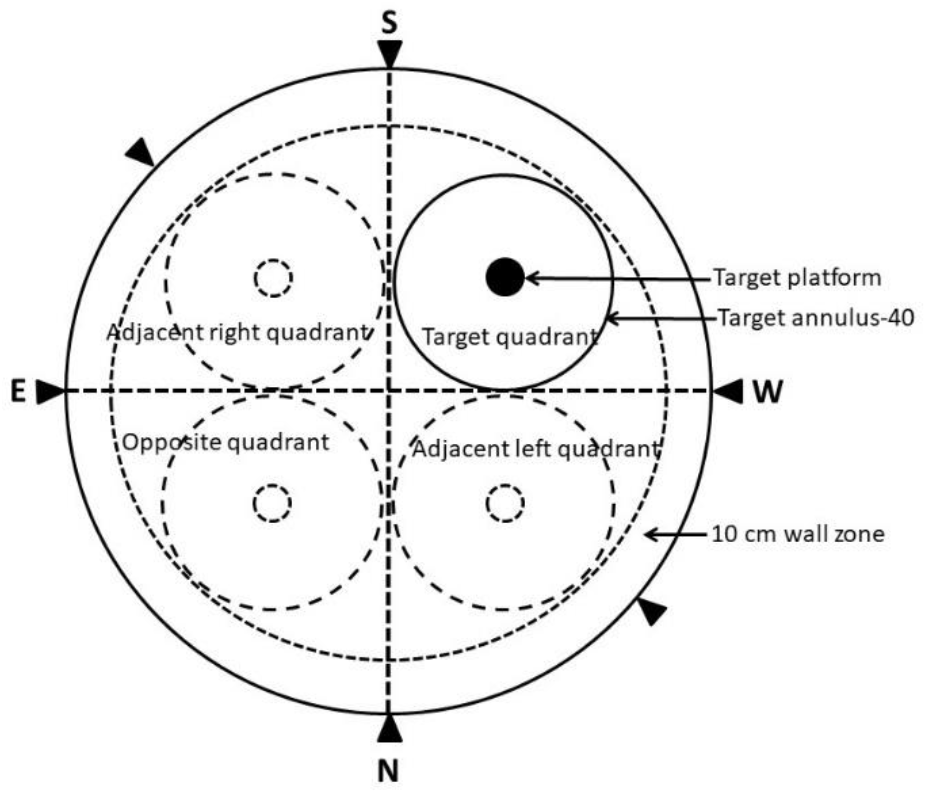
For the purpose of data analyses, the water maze was virtually divided into four quadrants and a 10-cm wide zone along the wall of the maze. The filled circle in the target (training) quadrant represents the target platform and the outer circle represents the annulus-40 in the target quadrant. The inner doted circles in the centers of non-target quadrants represent areas of hypothetical platforms and the outer doted circles represent hypothetical annuli-40.

## 3. Results

### 3.1. Effect of letrozole on estrous cyclicity

Exposure to both doses of letrozole (0.1 and 0.3 mg/kg s.c.) suspended estrous cyclicity after two weeks of the treatment.

### 3.2. Effect of letrozole on locomotor activity and locomotor habituation in open field

Locomotor habituation to open field is considered as a form of non-associative learning (Bolivar, 2009), and is strain dependent (Bolivar, 2009). It can be assessed in inter-session or intra-session paradigm (Bolivar, 2009). In the current study, we evaluated the effect of letrozole on inter-session habituation. For the total distance traveled (Figure 2A), two-way RM-ANOVA revealed significant effect of testing day [F(3,66)=3.941, p=0.012], non-significant effect of treatment, and significant testing day × treatment interaction [F(6,66)=2.276, p=0.047]. However, Tukey *post-hoc* comparison did not reveal significant differences among treatments-suggesting that overall locomotor performance was not affected by over 3 weeks of exposure to letrozole. To reveal in which of the tested group inter-session habituation occurred, for each group a within-subject one-way ANOVA was performed. The RM-ANOVA revealed significant differences among testing days only for letrozole 0.3 mg/kg group [F(3,31)=8.319, p<0.001], and Tukey pairwise comparison revealed significant decline of the distance traveled between day 1 and day 3 (p=0.001), and day 1 and day 4 (p=0.004). There was no significant decline of activity in mice in the control and the letrozole 0.1 mg/kg groups.

**Figure 2.**
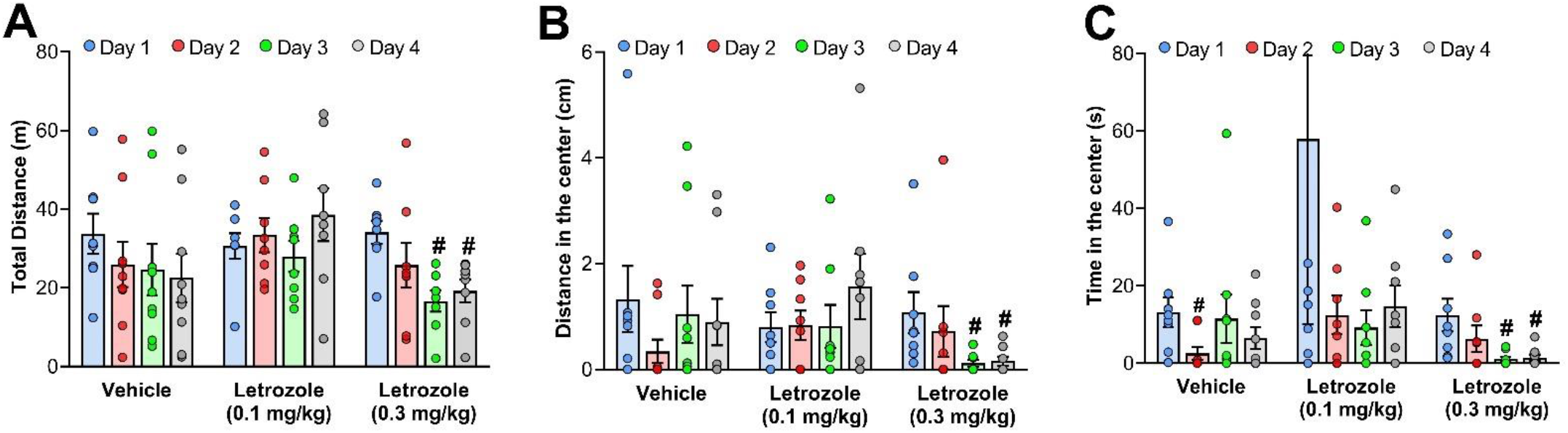
Locomotor habituation. Effect of letrozole on (A) total distance traveled, (B) distance in the center of arena, and (C) time in the center of arena, during 15-minute open field tests performed on 4 consecutive days. Testing was conducted during exposure to letrozole and following 3 weeks of prior exposure. Data points and error bars represent mean and SEM, respectively. Vehicle (n=9), Letrozole 0.1 mg/kg (n=8), Letrozole 0.3 mg/kg (n=8); exception: graph A for Day 1 for Vehicle represents mean and SEM only for 8 mice, because behavior of one mouse was disturbed during testing on this day; graph C for Day 1 for Letrozole 0.1 mg/kg does not show one mouse with the value 392 s, which is out of the scale range. (# p<0.05, compared with the 1^st^ day, according to within group analysis of RM-ANOVA or RM-ANOVA on Ranks, followed by Dunnet’s test).

To further characterize exploratory behavior, time and distance traveled in the center zone (20 cm × 20 cm) were evaluated (Figure 2B and C). For time spent in the center, two-way RM-ANOVA revealed non-significant effect of testing day, non-significant effect of treatment, and non-significant testing day × treatment interaction. For distance traveled in the center, two-way RM-ANOVA revealed non-significant effect of testing day, non-significant effect of treatment, and significant testing day × treatment interaction [F(6,66)=2.937, p=0.013]. An additional within group one-way RM-ANOVA analysis was performed for each group. For the control group, the one-way RM-ANOVA revealed significant decline of time and distance in the center only on the 2^nd^ day [time: RM-ANOVA on Ranks, χ2(3) = 13.329, p=0.004, followed by Tukey test, p<0.05; distance: RM-ANOVA on Ranks, χ^2^(3) = 11.351, p<0.001, followed by Tukey test p<0.05]. Thereafter, exploration of the center in the control group came back to the level of the 1^st^ day. In the letrozole 0.3 mg/kg group, there was significant decline of activity on the 3^rd^ and 4^th^day that was consistent with the decline of the total distance traveled [time: RM-ANOVA F(3,21)=5.021, p=0.009, followed by Tukey test, p<0.05; distance: RM-ANOVA on Ranks, χ^2^(3) = 16.500, p<0.001, followed by Tukey test p<0.05]. Letrozole 0.1 mg/kg group did not significantly alter activity over the course of testing. The high variability in this group on the first day of testing was related to the behavior of one mouse that spent almost all time (392 s) in the center. Results of the above analysis demonstrate that exploration of the open field center correlated with the overall exploratory activity and suggest lack of anxiogenic effect of exposure to letrozole on open field exploration.

### 3.3. Effect of letrozole on spontaneous alternation in Y-maze

Spontaneous alternation in Y-maze was performed 24 h after completing open field testing. At this time, mice were on letrozole treatment for 26 days. Spontaneous alternation in the Y-maze is based on natural propensity of rodents to explore a new place and is considered as a measure of spatial working memory. If memory is impaired, the number of alternations is decreased. On the other hand, if a treatment improves memory, the number of alternations is increased. None of the tested doses had a significant effect on spontaneous alternation (Tab. 1.)

**Table 1.** Spontaneous alternation in Y-maze; data represents mean and SEM.

|  | Vehicle | Letrozole<br>0.1 mg/kg | Letrozole<br>0.3 mg/kg |
| --- | --- | --- | --- |
| Percent of<br>alternations | 68.6 ± 2.5 | 75.0 ± 2.8 | 66.4 ± 4.1 |

### 3.4. Effect of letrozole on water maze performance

#### 3.4.1. Reference memory learning

Acquisition of the water maze task by female BALC/c mice was very slow; therefore, mice were trained in 5-day blocks interspaced with 2-days intervals until the average escape latency of any of the groups reached 20 s or less (Fig 3. A and B). Only the control group reached such criteria on the 15th day of the training. Two way repeated measure ANOVA conducted for a period of 15 days, revealed significant effect of training day [escape latency: F(14, 308) = 21.729, p < 0.01; distance: F(14, 308) = 12.933, p < 0.01], confirming progression of the learning process. However, effects of treatment, and treatment × training day interaction were not significantly different. The analysis was further narrowed to 5-day training blocks. Repeated measure ANOVA revealed significant effect of treatment for the 3rd block of training for escape latency [F(2,22)=3.529, p<0.05] and close to significance for distance travelled [F(2,22) = 3.004, p = 0.07]. Effects of treatment were not significant for the 1^st^ and the 2^nd^ training blocks. *Post-hoc* Tukey comparison for the 3^rd^ block revealed that treatment of mice with letrozole 0.1 mg/kg significantly impaired their learning, as their escape latency was significantly longer than that of control mice (p < 0.05). Effect of the training day was significant within the 1^st^ and the 2^nd^ block for escape latency [F(4,88)=7.640, p<0.01, and F(4,88)=2.667, p<0.05, for the 1^st^ and 2^nd^ block respectively], and within the 2^nd^ and 3^rd^ block for distance traveled [F(4,88)=5.093, p<0.001, and F(4,88)=5.034, p=0.001 for the 2^nd^ and 3^rd^ block respectively].Training day × treatment interaction were not significantly different within any of the 5-day blocks.

**Figure 3.**
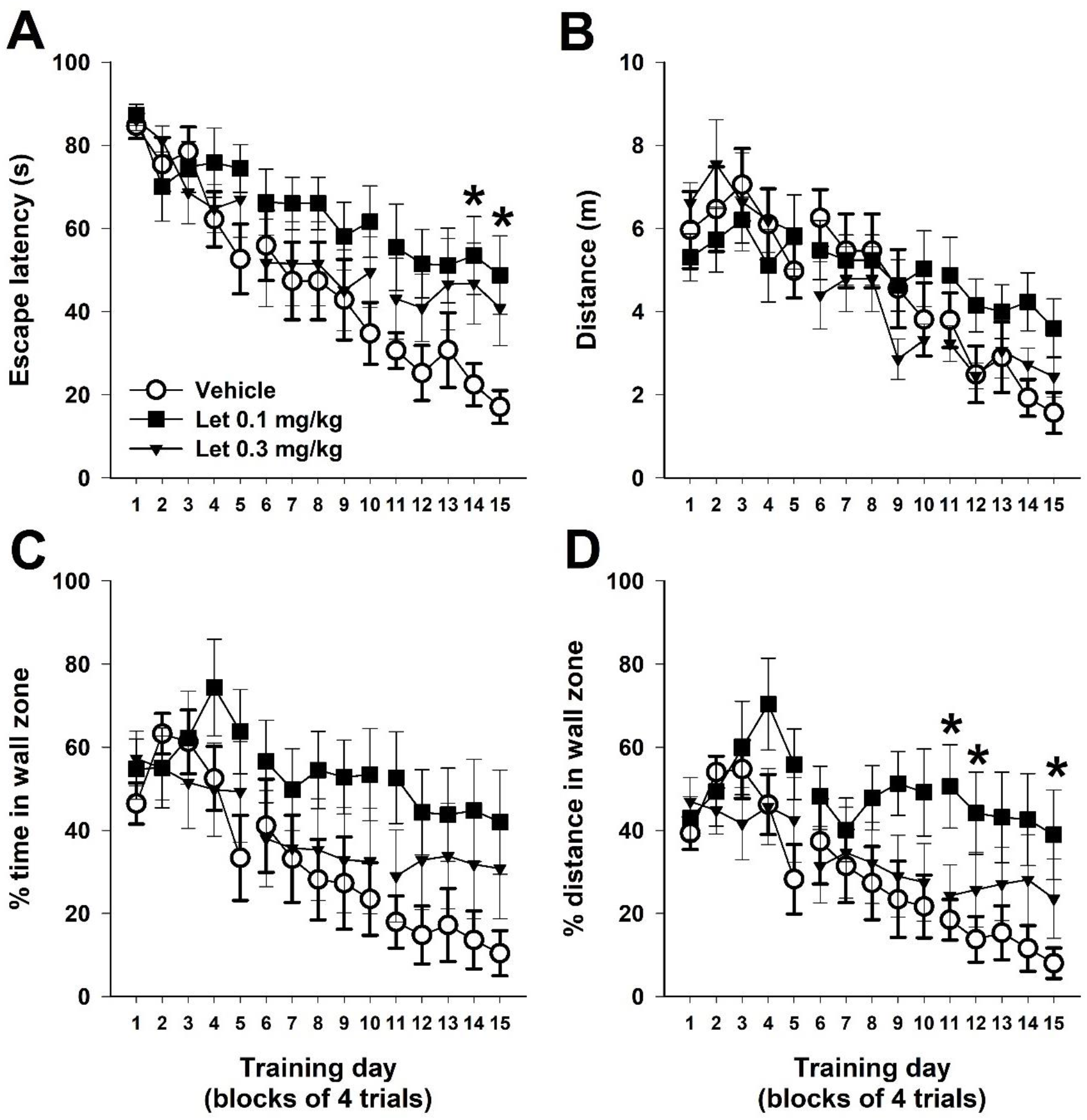
Learning performance of female BALB/c mice in the MWM. MWM training was conducted during exposure to letrozole and following 4 weeks of prior exposure. Graphs show (A) mean escape latency and (B) mean distance mice swam to find a submerged platform during 4 trials in each training day; and (C) mean percentage of the escape latency and (D) mean percentage of the distance that a mouse spent/traveled in 10 cm wide wall zone. Data points and error bars represent mean and SEM, respectively. Vehicle (n=9), Let 0.1 mg/kg (letrozole 0.1 mg/kg, n=8), Let 0.3 mg/kg (letrozole 0.3 mg/kg, n=8); * p < 0.05 vs Vehicle group, according to Tukey *post-hoc* test.

Mean swimming speed was not significantly different among treatments. RM-ANOVA reveal significant effect of the training day [F(14, 308)=4.768, p<0.001] and non-significant treatment × training day interaction, although close to significance, p=0.074. Narrowing analysis to 5-day training blocks did not reveal any significant effect of the treatment or the treatment × training day interaction. Interestingly the pattern of mean speed tended to have inverted-U shape, with the highest speed on the second week of the training (supplementary Fig. S1 A). In all tested group there were mice that had tendency for floating in wall zone, instead actively searching for the escape. There were 3 such mice in control group, 3 in letrozole 0.1 mg/kg group, and 2 in letrozole 0.3 mg/kg group. In the control group, all mice that initially floated actively escaped on the platform on the last week of the training. In contrast, in letrozole 0.1 mg/kg and in letrozole 0.3 mg/kg groups, most of these mice failed to mount the platform in 3 or 4 trials/4 trials on the last week of the training (supplementary Fig. S1 B).

To reveal the nature of the impairment, thigmotactic behavior was analyzed. The thigmotactic behavior was expressed as a percentage of the escape latency and the percentage of the distance that a mouse spent/traveled in the 10-cm wall zone in each trial. Thigmotactic behavior reflets anxiety (Simon et al., 1994; Treit and Fundytus, 1988) and is demonstrated as a tendency for swimming close to the wall around the perimeter of the pool (Dalm et al., 2000) during the first days of the training. It usually declines as animals learn how to find a hidden escape platform. In the current study, on the first training day, the % of time and distance traveled in the wall zone was similar among all experimental groups, and differences among groups became evident over the course of the training. Two-way RM-ANOVA revealed a significant main effect of training day for both % of time: [F(14, 308)=11.216, p<0.001] = 151.55, p < 0.0001] and % of distance traveled F(14, 308)=11.324, p<0.001] in the wall zone (Fig 3 C and D). The effect of the treatment was not significant. However, treatment × training day interaction was significant for both % of time [F(2,22) = 2.098, p = 0.01] and % of distance traveled [F(2,22) = 2.003, p = 0.002]. Mice exposed to letrozole 0.1 mg/kg had significantly increased % of time and % of distance in the wall zone during the last weeks of the training (Tukey *post hoc*, p < 0.05). Interestingly, the distance swum in the wall zone by mice treated with letrozole 0.1 mg/kg was significantly longer than that swum by control mice, although the total distance swum was not significantly longer. This result suggests that exposure to letrozole 0.1 mg/kg changed the navigation strategy relying on swimming to the platform using the shortest path through the center of the maze, to the strategy relaying on swimming to the platform around the perimeter of the maze.

To further determine the role of thigmotactic behavior in water maze impairment average escape latency was correlated with average percentage of time in wall zone (Fig 4). Escape latency to reach the platform was highly correlated with the percentage of time in wall zone, suggesting that thigmotaxis was a major factor contributing to the impaired performance (control group: R=0.986, p<0.001; letrozole 0.1 mg/kg: R= 0.915, p<0.005; letrozole 0.3 mg/kg: R= 0.986, p<0.001).

**Figure 4.**
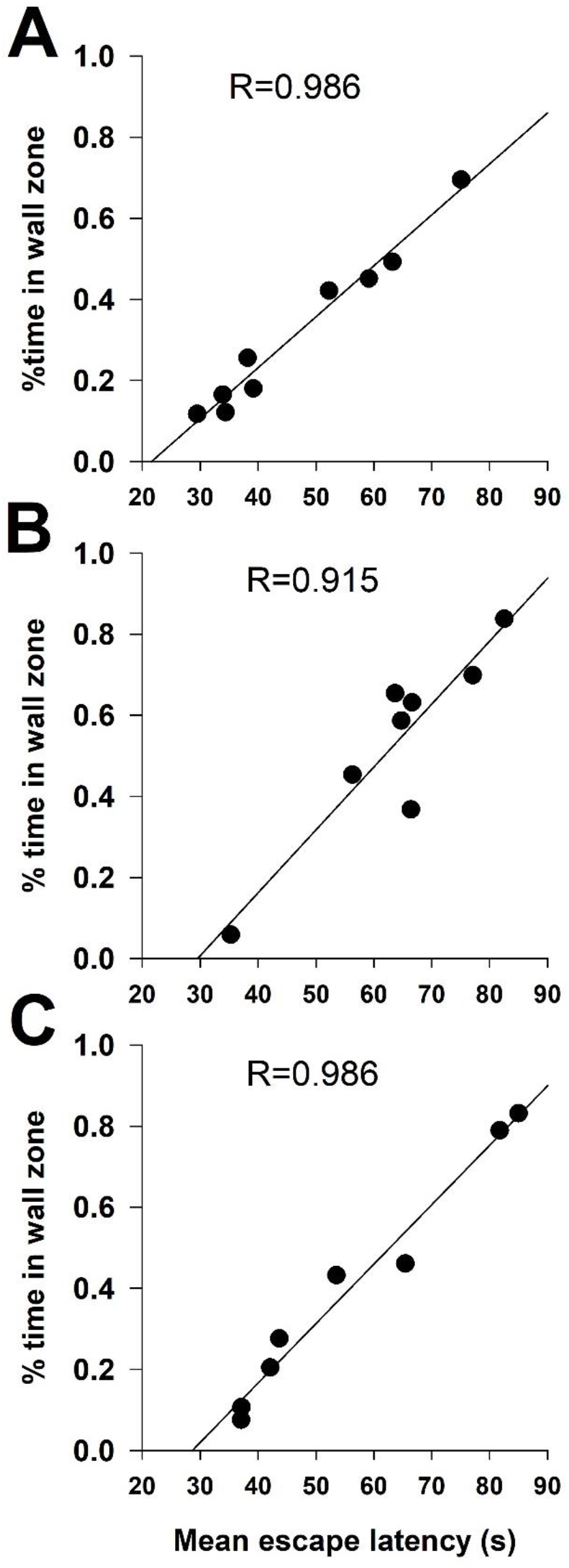
Correlation between mean escape latency and mean time spent in wall zone over period of 15-day reference memory training. (A) Vehicle, (B) Letrozole 0.1 mg/kg, and (C) Letrozole 0.3 mg/kg

#### 3.4.2. Reference memory probe test

A probe test was performed 72 hours after the last training trial. The platform was removed from the pool and mice were exposed to the water maze for 60 s. Motor performance measured as total distance traveled was not significantly different among groups (Tab. 2). Also, percentage of time and percentage of distance in 10 cm wall zone were not significantly different among groups, although there was strong trend for letrozole 0.1 mg/kg group to swim in the wall zone (Tab. 2).

**Table 2.** Reference memory probe test. Total distance swum during 60s probe test, and % of time and % of distance swum in 10 cm wide wall zone. Vehicle (n=9), Letrozole 0.1 mg/kg (n=8), Letrozole 0.3 mg/kg (n=8); Data represents mean and SEM

|  | Vehicle | Letrozole 0.1 mg/kg | Letrozole 0.3 mg/kg |
| --- | --- | --- | --- |
| Distance | 7.79 ± 0.68 | 4.98 ± 0.72 | 5.24 ± 1.08 |
| % of time wall zone | 22 ± 7 | 49 ± 10 | 35 ± 13 |
| % of distance wall zone | 15 ± 5 | 40 ± 9 | 31 ± 12 |

For cognitive performance measured as time and distance to reach the platform position, the longer time and distance indicate impairment. In the current study, if a mouse never crossed the platform position, a maximum time of 60 s and a total distance it swum were applied for the data analysis. For path efficiency to reach the platform, a value close to 1 indicates perfect navigation and a value close to 0 indicates severe impairment. For persistency for searching in a target platform position, values higher than a quarter of the total time or distance indicate better memory, and values lower indicate impairment.

Time to the first platform position entry was significantly longer for mice treated with letrozole 0.1 mg/kg or letrozole 0.3 mg/kg than for control mice [one way ANOVA F(2,24) = 8.222, p=0.002, followed by Tukey *post hoc* test, p = 0.002 and p = 0.024 for letrozole 0.1 mg/kg and letrozole 0.3 mg/kg, respectively] (Fig. 5 A), while distance traveled was significantly longer only for mice treated with letrozole 0.1 mg/kg group [Kruskal-Wallis ANOVA on Ranks, H = 8.697 with 2 degrees of freedom, p=0.013, followed by Dunn’s Method, p<0.05) (Fig 5 B). Navigation of control mice was very efficient with path efficiency scoring above 0.7, while path efficiency of mice treated with letrozole 0.1 mg/kg was below 0.2 and significantly lower compared to control mice [one way ANOVA F(2,24) = 7.161, p = 0.004, followed by Tukey *post hoc* test, p = 0.003]. Path efficiency of mice treated with letrozole 0.3 mg/kg was intermediate (0.5) and was not significantly different from that of mice treated with vehicle (control) or letrozole 0.1 mg/kg (Fig 5. C). These data indicate significant impairment of navigation in mice treated with letrozole 0.1 mg/kg.

**Figure 5.**
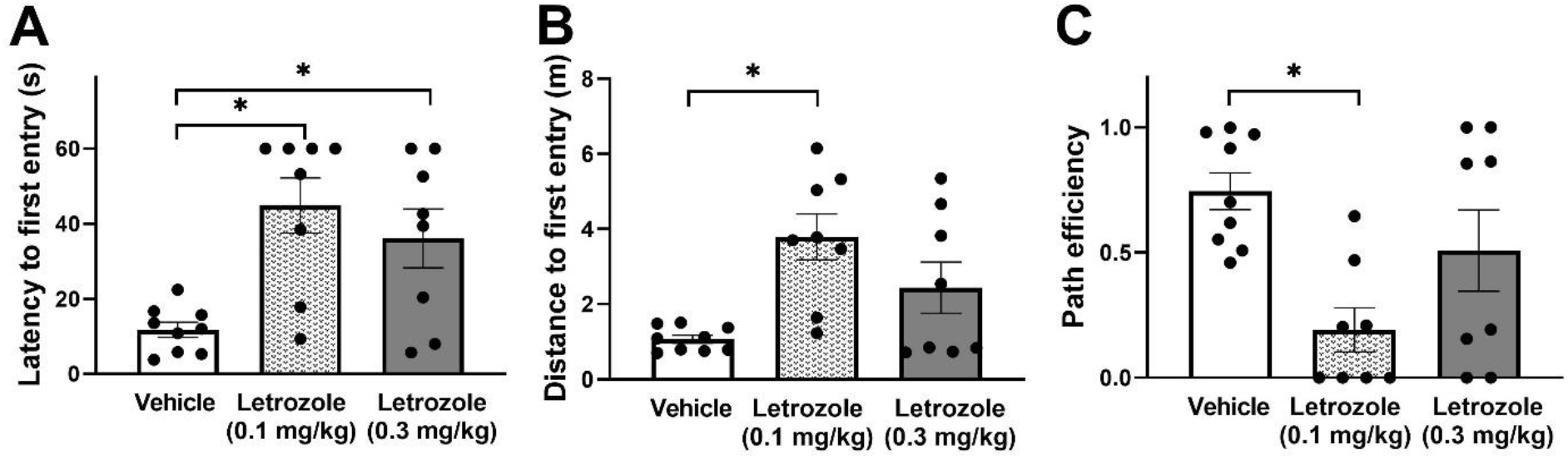
Performance of mice in a probe test performed 72 h after the last training trial, measured as (A) time, (B) distance, and (C) path efficiency for the first entry to the target platform position. Data points and error bars represent mean and SEM, respectively. Vehicle (n=9), Letrozole 0.1 mg/kg (n=8), Letrozole 0.3 mg/kg (n=8) (* p<0.05, one-way ANOVA followed by Tukey *post-hoc*).

Persistence for searching in the target platform position was evaluated between groups as well as within each group on three levels of accuracy: quadrant, annulus-40, and platform area crossings. Importance of the of evaluation of preference on the level of anululs-40 relies on the fact that this area does not include wall zone and evaluates only active search for the target platform, thereby, excluding mice that passively floated in the wall zone.

Between groups comparison showed that mice treated with letrozole 0.1 mg/kg group swam significantly shorter distance in the target quadrant than control mice [one way ANOVA F(2,24)=3.956 p=0.034, followed by Tukey *post-hoc* test, p=0.03] (Fig. 6 B), but time spent in the target quadrant was not significantly different among groups (Fig. 6 A). In the area of closer proximity to the target platform center as annulus-40, mice treated with letrozole 0.1 mg/kg spent shorter time [one way ANOVA F(2,24)= 3.576, p=0.045, followed by Tukey *post hoc* test, p=0.041] and swam shorter distance than control mice [one way ANOVA F(2,24)= 5.029, p=0.016, followed by Tukey *post hoc* test, p=0.013] (Fig 6 C and D). Number of target platform area crossings was also significantly lower for mice treated with letrozole 0.1 mg/kg than control mice [one way ANOVA on Ranks, H = 9.073 with 2 degrees of freedom p=0.011, followed by Dunn’s Method, p<0.05) (Fig 6 C).

**Figure 6.**
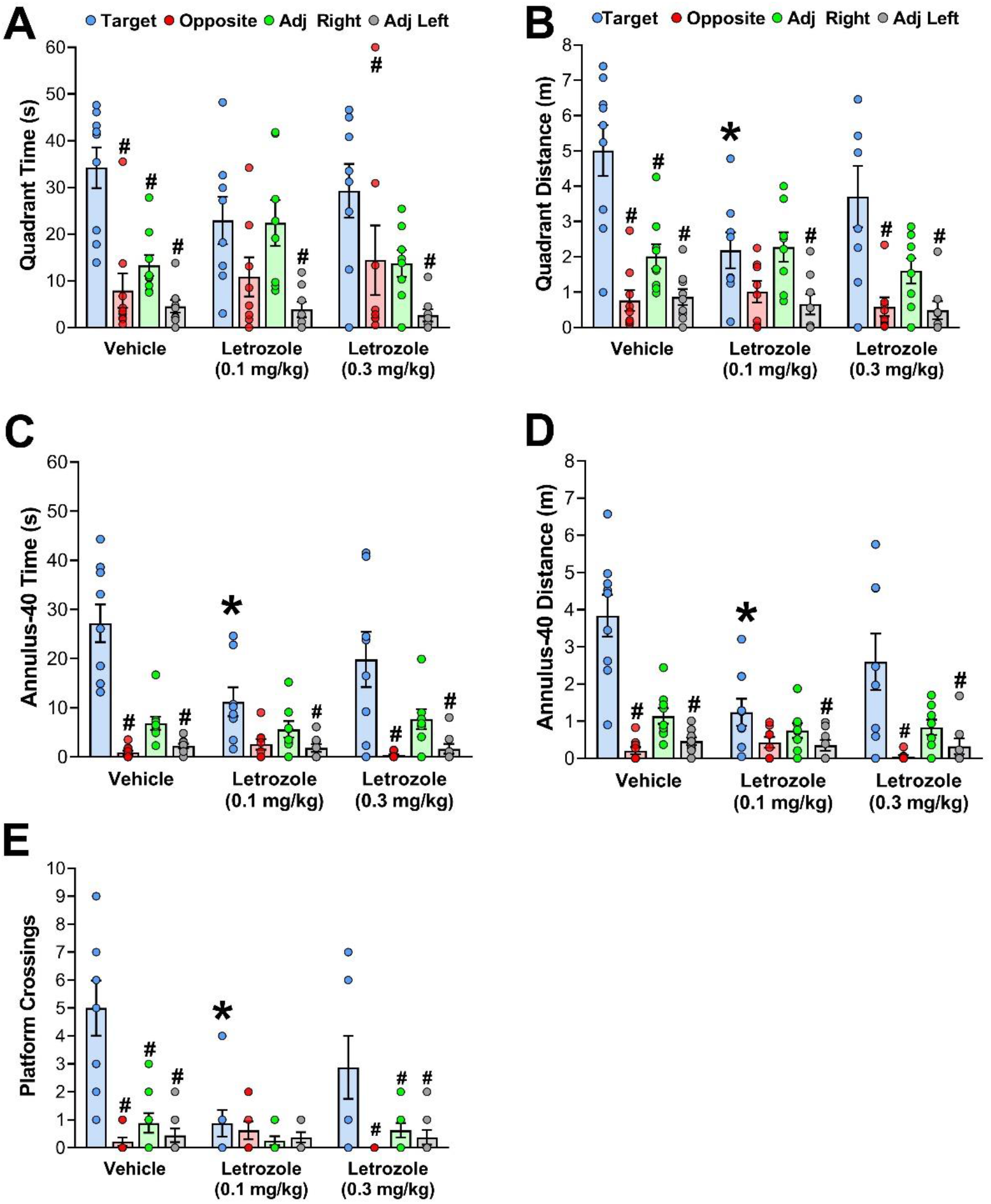
Performance of mice in a probe test performed 72 hours after the last training trial. The performance was analyzed on the level of quadrants: (A) time and (B) distance; annulus-40: (C) time and (D) distance; and platform size areas: (C) crossings. For among groups comparison one-way ANOVA was conducted for time and distance in the target quadrant, for time and distance in the target anulus-40, and for number of crossings of the target platform position Data points and error bars represent mean and SEM, respectively. Vehicle (n=9), Letrozole 0.1 (n=8), Letrozole 0.3 (n=8) (* p<0.05, one-way ANOVA followed by Tukey *post hoc*). For within group comparison, time spent in, and % of distance swum in target quadrant, or target annulus-40, was compared with time and distance in non-target quadrants, or annuli-40 virtually positioned in the center of non-target quadrants; target platform position crossings was compared to crossing of platform size areas virtually positioned in centers of non-target quadrants (# p<0.05, RM-ANOVA or RM-ANOVA on Ranks, followed by Dunnet’s test)

For the within-group comparison, time spent, distance swum, and crossings of the target quadrant, target annulus-40, and target platform area were compared to equivalent areas located in centers of other quadrants. Control mice showed significant preference to the target quadrant vs opposite and adjacent quadrants both for the time [RM-ANOVA F(3,35)=13.452, p<0.001, followed by Dunnet’s test p<0.001] and distance [RM-ANOVA on Ranks, χ^2^(3) = 20.600, p<0.001, followed by Dunnett’s test, p<0.05) (Fig.6. A and B). Mice treated with letrozole 0.3 mg/kg also showed significant preference to the target quadrant for time and distance [RM-ANOVA on Ranks, χ^2^(3) = 10.263, p=0.016, followed by Dunnett’s test p<0.05; and χ^2^(3) = 11.526, p= 0.009, followed by Dunnett’s test p<0.05, for time and distance, respectively] (Fig 6 A and B). For mice treated with letrozole 0.1 mg/kg, although ANOVA was significant, there was no significant preference to the target quadrant vs the opposite quadrant, both for time [repeated measure ANOVA F(3,35)= 3.704, p=0.028] and distance [RM-ANOVA on Ranks, χ^2^(3) = 9.835, p=0.016] (Fig 6 A and B).

On the level of annulus-40 (Fig 6 C and D), control mice showed significant preference to target area versus area in the center of the opposite quadrant \[RM-ANOVA on Ranks, χ^2^(3) = 25.705, p<0.001, followed by Dunnett’s test p<0.05; and χ^2^(3) = 22.483, p<0.001, followed by Dunnett’s test p<0.05, for time and distance, respectively]. Similar results were observed with mice treated with letrozole 0.3 mg/kg [RM-ANOVA on Ranks, χ^2^(3) = 16.324, p<0.001, followed by Dunnett’s test p<0.05; and χ^2^(3) = 16.324, p<0.001, followed by Dunnett’s test p<0.05, for time and distance, respectively]. For mice treated with letrozole 0.1 mg/kg, although ANOVA was significant, there was no significant preference to the target quadrant vs the opposite quadrant [RM-ANOVA on Ranks, χ^2^(3) = 8.919, p=0.03. and χ^2^(3) = 8.595, p=0.035, for time and distance, respectively].

For platform crossing (Fig 6 F), significant preference to target area versus equivalent area in the center of the opposite quadrant was evident in the group of control mice [RM-ANOVA on Ranks, χ^2^(3) = 17.878, p<0.001, followed by Dunnett’s test p<0.05] and the group of mice treated with letrozole 0.3 mg/kg [repeated measure ANOVA F(3,35)= 5.007, p= 0.009]. There was not significant deference among positions for mice treated with letrozole 0.1 mg/kg.

#### 3.4.3. Reversal training

To evaluate if exposure to letrozole affected learning flexibility, mice were subjected to reversal learning 24 h after completing the probe test. For this reversal training, the platform was moved to the center of the opposite quadrant and mice were trained for 5 consecutive days to find it. Mice learnt the new platform position relatively quickly in comparison with the first platform position. Four of five letrozole mice that were not able to actively search for the platform during the reference memory training finally were successful in navigating toward it on the 5^th^ day of the reversal training. Two-way RM-ANOVA revealed significant effect of training day [escape latency: F(4,88)=26.259, p<0.01; distance: F(4,88)=37.607, p<0.01], but not of treatment, or treatment × training day interaction. For percentage of time and distance in the wall zone two-way RM-ANOVA revealed significant effect of training day [escape latency: F(4,88)=9.6, p<0.01; distance: F(4,88)=6.08, p<0.022], but not of treatment, or treatment × training day interaction.

**Figure 7.**
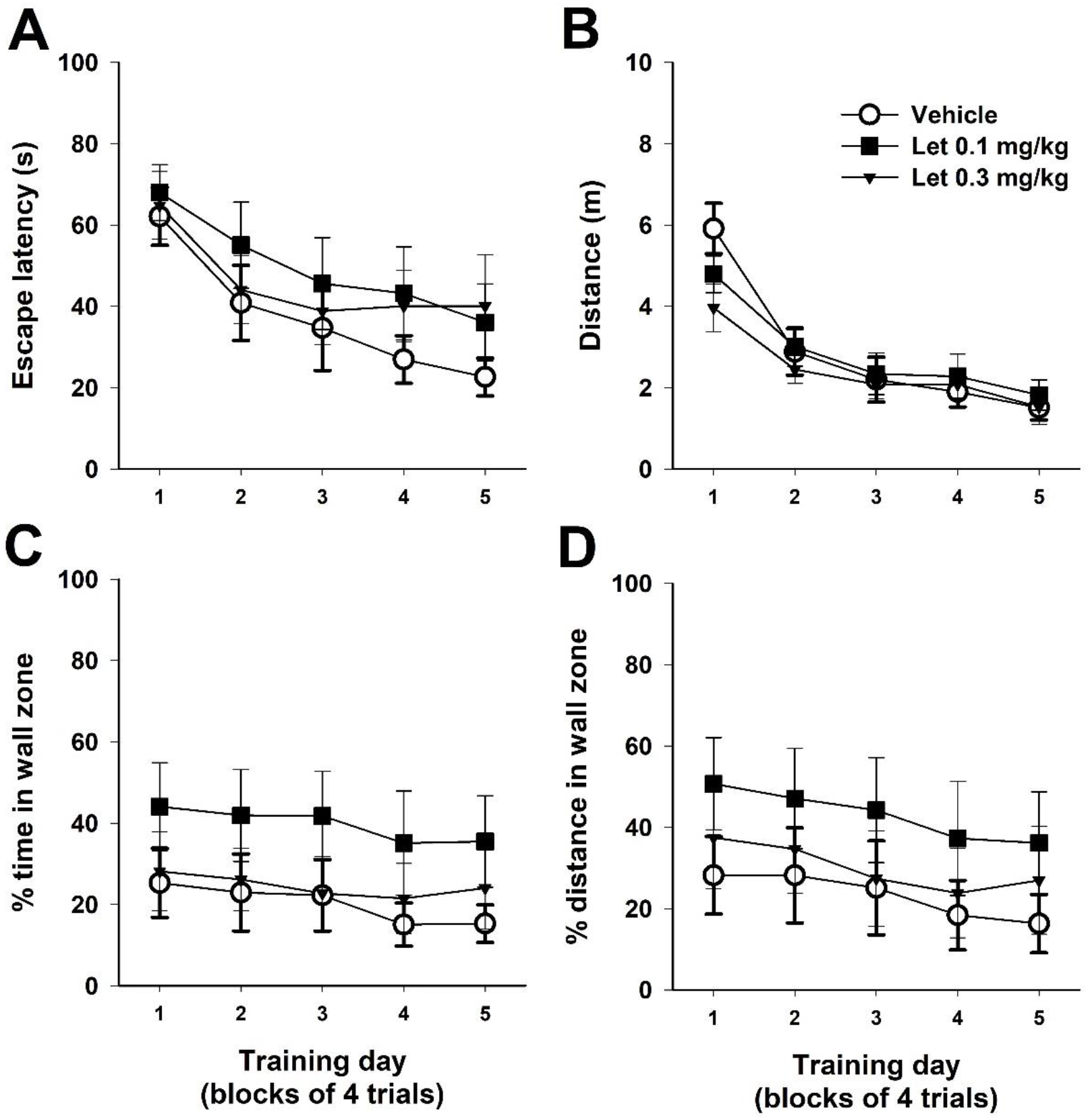
Reversal learning performance of female BALB/c mice in the Morris water maze. Reversal training was conducted during exposure to letrozole and following 7 weeks of prior exposure. Graphs show (A) mean escape latency and (B) mean distance mice swam during 4 trials in each training day to find a submerged platform; and (C) percentage of the escape latency and (D) percentage of the distance that a mouse spent/traveled in 10 cm wall zone. Data points and error bars represent mean and SEM, respectively. Vehicle (n=9), Let 0.1 mg/kg (letrozole 0.1 mg/kg, n=8), Let 0.3 mg/kg (letrozole 0.3 mg/kg, n=8)

#### 3.4.4. Post-reversal probe test

A post-reversal learning probe test was performed 72 h after completing the last trial of the 5-day training. This time, mice were allowed to swim 90 s. Mice that were not able to successfully find the platform were not included for the analysis. Two mice that found the platform only on the two last reversal training trials were excluded from the letrozole 0.1 m/kg group, and one mouse that had never found the platform was excluded from the letrozole 0.3 mg/kg group. All the excluded mice spent most of the reversal training time in the wall zone.

During the probe test, total distance traveled and percentage of time and percentage of distance in 10 cm wall zone were not significantly different among groups (Tab 3). Cognitive performance measured as time to the first platform position entry and path efficiency was not significantly different among group. Time spent or distance swum in target quadrant, target annul-40, and number of crossings of the target platform position were also not significantly different among groups.

**Table 3.** Reference memory probe test. Total distance swum during 60s probe test, and % of time and % of distance swum in 10 cm wide wall zone. Vehicle (n=9), Letrozole 0.1 mg/kg (n=6), Letrozole 0.3 mg/kg (n=6); Data represents mean and SEM

|  | Vehicle | Letrozole 0.1 mg/kg | Letrozole 0.3 mg/kg |
| --- | --- | --- | --- |
| Distance | 10.2± 0.9 | 8.21± 1.23 | 8.26 ± 1.72 |
| % of time wall zone | 31.4± 2.5 | 37.6 ± 12.4 | 29.2± 12.8 |
| % of distance wall zone | 25.4 ± 9.0 | 31.0 ±11 | 20.2 ± 9.0 |

To reveal preference to the new platform position versus old position, a within-group comparison was performed. On the level of quadrants, control mice showed significant preference to the target quadrant vs opposite and all adjacent quadrants both for time [RM-ANOVA F(3,35)= 26.918, p<0.001, followed by Dunnet’s test p<0.001] and % of distance [RM-ANOVA F(3,35)= 26.597, p<0.001, followed by Dunnet’s test p<0.001]. Mice treated with letrozole 0.1 mg/kg also showed significant preference to the target quadrant vs opposite for time [RM-ANOVA F(3,23)= 12.519, p<0.001, followed by Dunnet’s test p<0.001], and % of distance [RM-ANOVA on Ranks, χ^2^(3) = 13.862, p=0.003, followed by Dunnett’s test p<0.001]. Mice treated with letrozole 0.3 mg/kg showed significant preference to the target quadrant vs opposite only for time [RM-ANOVA F(3,27)= 7.675, p=0.001, followed by Dunnet’s test p<0.005] but not for % of distance [RM-ANOVA F(3,27)= 3.883, p=0.027, followed by Dunnet’s test] (Fig 8 A and B).

**Fig. 8.**
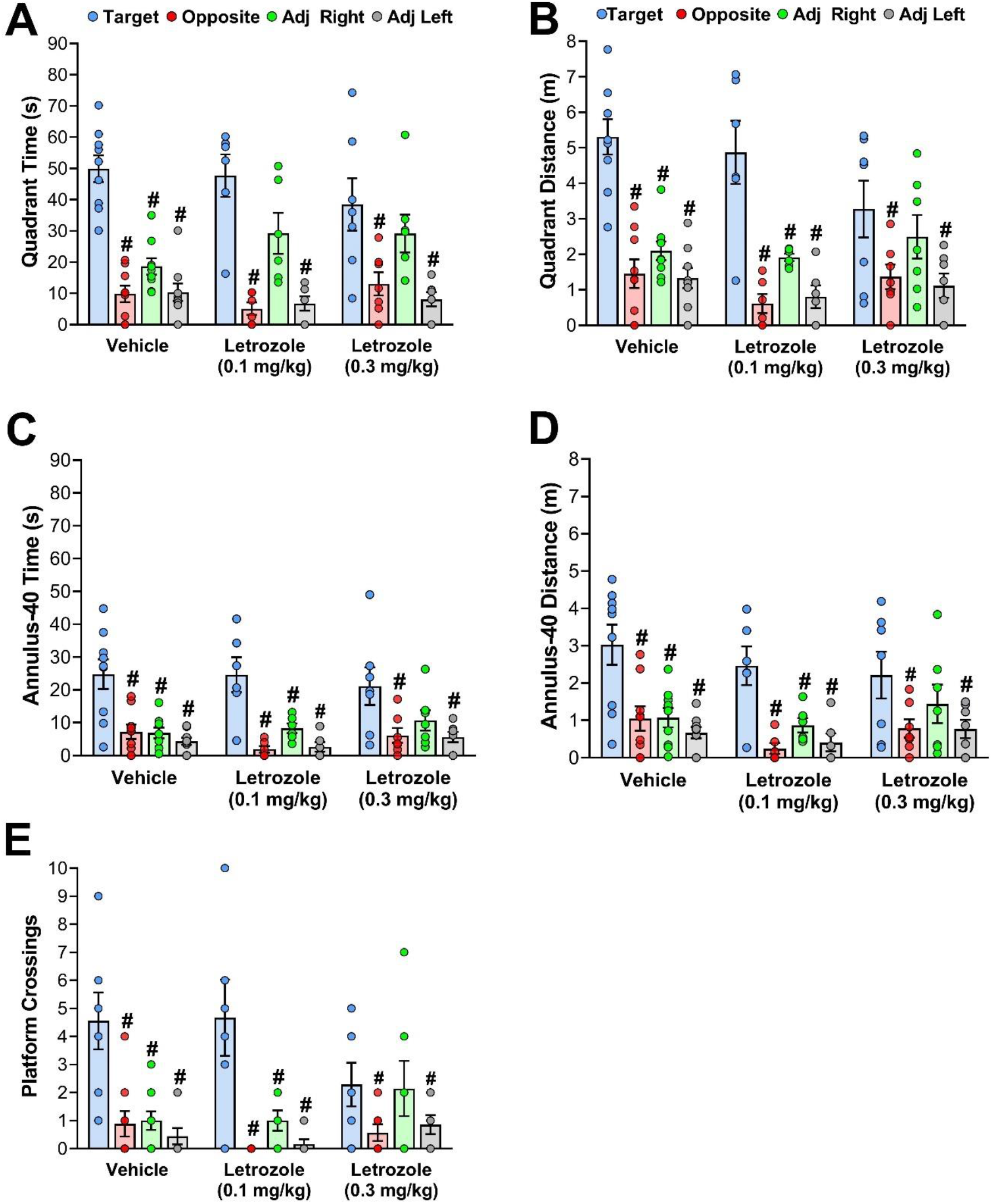
Performance of mice in a post-reversal probe test performed 72 hours after the last training trial. The performance was analyzed on the level of quadrants: (A) time and (B) distance; annulus-40: (C) time and (D) distance; and platform size areas: (C) crossings. For within group comparison time spent in, distance swum in, and platform crossings of areas related to the target platform positions was compared versus time spent in, % of distance swum in, and platform crossings in non-target quadrants and areas virtually positioned in its centers Data points and error bars represent mean and SEM, respectively. Vehicle (n=9), Letrozole 0.1 mg/kg (n=6), Letrozole 0.3 mg/kg (n=7) (# p<0.05, repeated measure ANOVA or Friedman Repeated Measures Analysis of Variance on Ranks followed by Dunnet’s test)

On the level of annulus-40, control mice showed significant preference to the target annulus-40 vs opposite and all adjacent quadrants both for the time [RM-ANOVA F(3,35)= 18.158, p<0.001, followed by Dunnet’s test p<0.05] and distance [RM-ANOVA on Ranks, χ^2^(3) = 26.333, p<0.001, followed by Dunnett’s test p<0.001]. Mice treated with letrozole 0.1 mg/kg also showed significant preference to the target annulus-40 for time [RM-ANOVA F(3,23)= 19.741, p<0.001, followed by Dunnet’s test p<0.001] and distance [RM-ANOVA on Ranks, χ^2^(3) = 14.895, p= 0.002, followed by Dunnett’s test p<0.05]. Likewise, mice treated with letrozole 0.3 mg/kg showed significant preference to the target annulus-40 for time [RM-ANOVA on Ranks, χ^2^(3) =9.957, p=0.019, followed by Dunnett’s test p<0.001] and distance [RM-ANOVA F(3,27)= 3.747, p= 0.030] (Fig 8 C and D).

On the level of platform area crossings vs equivalent area in centers of the opposite quadrant, control mice showed significant preference to the target platform position [RM-ANOVA F(3,35)= 12.920, p<0.001, followed by Dunnet’s test p<0.001]. Mice treated with letrozole 0.1 mg/kg also showed preference to the target platform position [RM-ANOVA F(3,23)= 11.452, p<0.001, followed by Dunnet’s test p<0.001], whereas mice treated with letrozole 0.3 mg/kg did not (Fig.8 E).

The above results indicate that mice treated with letrozole 0.1 mg/kg were finally able to successfully navigate in the water maze. The tested doses of letrozole did not induce a significant interference with the previous platform position suggesting no effect of treatment on learning flexibility. Surprisingly, mice treated with letrozole 0.3 mg did not show significant preference to the target platform measured as of distance on the level of quadrant and annulus-40, as well as platform crossings, suggesting that, in certain contexts, the higher dose of letrozole can disturb learning/memory process.

#### 3.4.5. Reference memory training in a new room

Twenty-four hours after completing the reversal memory probe test, mice were brought to a new room and tested in a water maze with the same dimensions and similar lighting conditions on the water surface as the previous one. The training was conducted in two 4-day blocks interspaced by 72 h break. Repeated measure ANOVA did not reveal significant effect of the treatment or treatment × training day interaction, but it revealed significant effect of the training day [escape latency: F(7,154)=5.453, p<0.01; distance: F(7,154)=10.472, p<0.01]. The performance of all mice was not as good as expected for water maze-experienced mice. All mice that showed highly thigmotactic behavior during the first reference memory training expressed it again and were not able to escape on the submerge platform. Interestingly, before the training in a new room the last time control mice expressed such behavior on the second week of the first reference memory training. The learning curve was flat on the first week of the training, with control mice improving their performance after the 72-h brake, on the 5th day of the training. On this day, the performance of control mice was significantly different than that of mice treated with letrozole 0.1 mg/kg or 0.3 mg/kg, as revealed by one-way ANOVA. However, control mice did not express any further improvement on the following days. There might be several reasons for the poor performance, (i) the water temperature was kept at 22 °C, and this low temperature might have inhibited habituation to the novel testing environment, (ii) the new testing environment could have been too anxiogenic for mice, (iii) mice could have lost motivation for the escape after prolonged testing (supplementary Fig S2).

#### 3.4.6. Post-MWM open field

Post-MWM testing in a large open field (120 cm × 120 cm) did not reveal significant differences in the exploratory behavior measured as a total distance traveled and the distance traveled in center of the open field (defined as an area outside the 20 cm zone around the wall) (supplementary Fig. S3). This result suggests the letrozole did not alter anxiety-like behavior in the open field.

To assess if the propensity for swimming in the MWM wall zone can be predicted based on the anxiety-like behavior of an individual mouse in the open field, average percentage of distance in the MWM wall zone during the 3^rd^ week of reference memory training was plotted against percentage of distance in the center of the open field during the first 20 min of the test (supplementary Fig. S4). The 3^rd^ week of the reference memory training was selected for analysis because there was significant difference among groups. Pearson’s correlation analysis did not reveal any significant relationship between both measures, suggesting that anxiety like-behavior of female BALB/c mice in the MWM is independent form their anxiety-like behavior on dry land.

## 5. Discussion

The current study showed that letrozole dose-dependently modulated behavioral response and that its effect was task-dependent. Each of tested doses had different effect in locomotor habituation learning task and in water maze learning task.

Letrozole at the dose of 0.3 mg/kg facilitated locomotor habituation while the dose of 0.1 mg/kg had tendency to disrupt it. As locomotor habituation is considered a form of non-associative learning, the action of letrozole at the dose of 0.3 mg/kg may be recognized as pro-cognitive. In fact, certain doses of letrozole have been reported to augment cognition in rodents (Aydin et al., 2008) and monkeys (Gervais et al., 2019). Both intra- and intersession habituation in different mouse strains has been reported to be sensitive to genetic manipulations and drug treatment (Leussis and Bolivar, 2006). However, to date, there have not been any publication describing effects of letrozole or sex hormones as e.g., estrogen or testosterone, on locomotor habituation. Although locomotor habituation was not significant in control mice, there was an observable decline of activity that could possibly have reached statistical significance after additional habituation sessions. Previous study of inter-session locomotor habituation in BALB/c mice did not find any decline of locomotor activity regardless of whether the study used short, 5-min tests (Bolivar et al., 2000) or long, 2-hour tests (Isles et al., 2004) on three consecutive days. The observed decline in our study could be an effect of different experimental protocols, involving in our study everyday injections, and prior habituation to the testing room and to the testing procedure, as well as longer testing time.

Letrozole 0.3 mg/kg did not significantly affect reference learning and memory in the MWM, while the lower dose of 0.1 mg/kg significantly impaired it. The impairment was more pronounced during the probe test than during the acquisition, a finding that may be related to: (i) the 72-h break enhancing differences among groups, and/or (ii) the nature of the probe test that is based on persistence in searching for the platform position and gives premium to animals that actively search for it. Impairment induced by the letrozole dose of 0.1 mg/kg is in agreement with previously reported impairment of water maze reference memory learning caused by similar dose, 0.16 mg/kg, injected intraperitoneally (i.p.) in female C57Bl mice (Liu et al., 2019). Similarly to an effect in female mice, i.p. administration of letrozole at a dose of 0.08 mg/kg has been reported to impair water maze learning in male C57Bl mice (Zhao et al., 2018). The dose of 0.3 mg/kg tested in the present study seems to be above the dose range that can generate impairment of water maze learning. A letrozole dose as high as 0.4 mg/kg administered s.c. (Vierk et al., 2015), and 2.5 mg/kg administered orally (Meng et al., 2011) were reported to have no effect on water maze learning/memory in C57Bl/6J mice. In addition, in Sprague-Dawley rats, the doses of 1 mg/kg, and 2.5 mg/kg, administered orally, were reported to improve reference memory learning in water maze (Aydin et al., 2008), and working memory in cross-arms maze (Alejandre-Gomez et al., 2007), respectively. Inconsistent effects of letrozole on learning and memory in rats and mice may be unique to these species. Alternatively, they may represent off-target effects of letrozole because, at least based on allometric conversion of doses, the letrozole doses studied in rats convert to human equivalent doses that are well above those approved for breast cancer treatment.

The observed impairment of spatial reference learning and memory in BALB/c mice exposed to letrozole dose 0.1 mg/kg could be an effect of increased anxiety rather than solely impaired navigation. There are few facts supporting this hypothesis. We observed many unsuccessful attempts to escape on the platform because mice did not swim far enough from the wall of the tank and came back to the wall zone. Interestingly, all highly anxious control mice were able to successfully escape on the platform on the last two days of the reference memory training, while 4 of 5 highly anxious mice exposed to letrozole were able to successfully escape only on the 5th day of the reversal training. The results of the reversal test proved that highly anxious mice can learn the water maze task, but they require a longer training to initiate searching in the center of the maze. Occurrence of highly anxious BALB/c mice that predominantly float has been reported by others (Francis et al., 1995; Johns et al., 2021; Philpot et al., 2016; Wahlsten et al., 2005). In some studies, such mice were excluded from testing. One study (Philpot et al., 2016) described removal from the study of 13 of 38 females, and 1 of 18 males, and other (Johns et al., 2021) described removal of 16 of 50 females. In those studies, water was kept at 24 °C. It seems that our approach with gradual decrease of water temperature and prolonged period of training facilitated performance of BALB/c mice in the water maze task, allowing for the successful task acquisition of all control mice.

BALB/c mice are highly emotional. Their emotionality is manifested by higher corticosterone surge during response to novelty than in C57BL/6 strain (Brinks et al., 2007). Despite strong neophobic reaction to novelty BALB/c mice have been characterized as having adaptive anxiety phenotype that develops habituation towards a novel stimulus over time (Salomons et al., 2012). They are also considered as superior learners in dry mazes (Brinks et al., 2007). The result of the current study suggests that letrozole 0.1 mg/kg could disrupt habituation process and delay learning in water maze. Evidence of importance of habituation process to novel environment was the fact that improvement of performance of highly anxious mice was transient. After moving to a new room, they expressed again anxiety behavior, regardless of the treatment, and leaned toward the wall zone. Similar phenomenon was described by (Huang et al., 2012a), where already pre-trained mice impaired their performance in the same water maze after increasing light intensity. This evidence suggests that for BALC/c mice, habituation to the testing condition is critical for successful performance in water maze. Letrozole-induced changes in emotionality could be a key factor leading to delay in acquisition of water maze task, as aromatase is mainly expressed within hypothalamic and limbic brain regions, including hippocampus and amygdala (Roselli, 2007) that control emotionality and response to stress (Barkus et al., 2010)

To explain difference between doses of letrozole, as well differences between mice and rats, the levels of testosterone have also to be considered (Park et al., 2009). Most of the publications explain the behavioral changes seen in letrozole-treated animals as the result of the lack of E2. However, estrogens and androgens can compensate for each other’s actions, and the behavioral changes induced by letrozole could be due not only to the lack of estrogen but the elevated levels of androgens. It is recognized by now that the concept of “male” and “female” hormones is an oversimplification of a complex developmental and biological network of steroid actions that directly impacts several organs. To complicate the situation further, acute stress, which increases the release of corticosterone from the adrenal gland, significantly enhances aromatase activity in the preoptic area of quails (Brooks et al., 2012; Dickens et al., 2014) and rapidly increases aromatase expression and E2 levels in the paraventricular nucleus of proestrus and ovariectomized female rats (Lu et al., 2019) Although these studies focused on the regulatory effects of aromatase in the hypothalamus, it remains unknown whether similar effects develop in the hippocampus or in amygdala. If stress increases aromatase activity in the hippocampus and amygdala, then, the elevated levels of testosterone together with corticosterone may be another confounding factor playing a role in the heterogeneity of the behavioral findings reported by different research groups studying letrozole-treated rodents.

In conclusion, the result of the current study demonstrate that BALB/c female mice are suitable for studying detrimental effects of letrozole on spatial learning/memory in water maze, although they require significantly higher effort than C57BL mice to be trained. The water maze also emerges as a relevant test to captured letrozole-induced cognitive impairment. Although the current study does not rule out direct negative effect of letrozole on spatial navigation, disturbance of navigation due to increased anxiety needs to be also considered when interpreting effects of letrozole on the water maze performance.

## Acknowledgements

The authors are indebted to Dr. Edna Pareira for her assistance in planning and conducting experiments, when she worked at the University of Maryland, School of Medicine.

We acknowledge the support of the University of Maryland, Institute for Clinical & Translational Research (ICTR) and the National Center for Advancing Translational Sciences (NCATS) CTSA grant number 1UL1TR003098.

## Author contributions

Jacek Mamczarz - Methodology, Investigation, Formal analysis, Visualization, Writing - Original Draft, Writing - Review & Editing

Malcolm Lane - Investigation

Istvan Merchenthaler - Funding acquisition, Conceptualization, Writing - Review & Editing

## SUPPLEMENTARY MATERIALS

**Figure S1.**
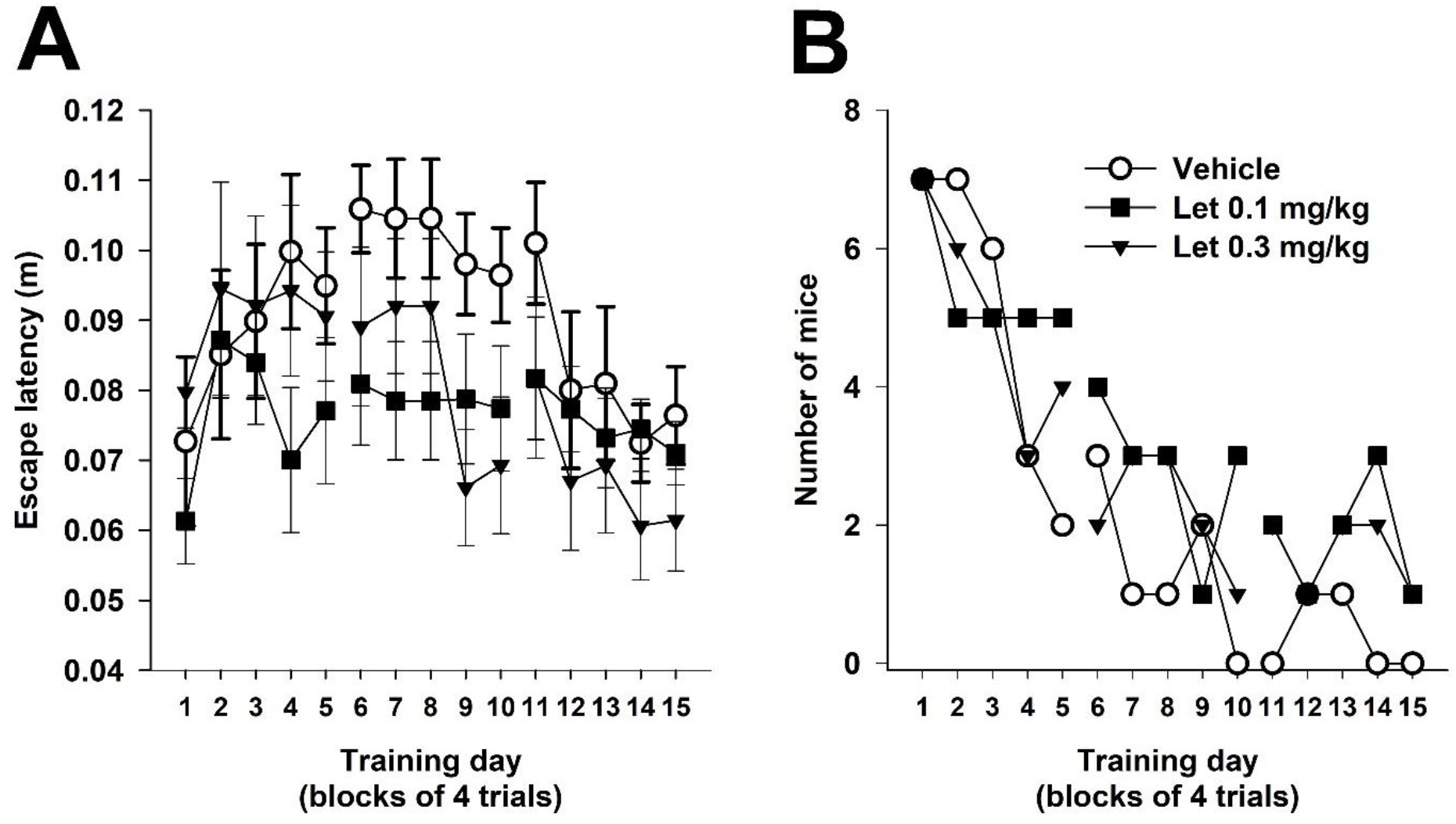
(A) Mean speed during reference memory training; data points and error bars represent mean and SEM, respectively. (B) Number of mice that failed to mount on the platform in 3-4 trials per 4 trials during reference memory training. Data points represent number of mice. Vehicle (n=9), Letrozole 0.1 mg/kg (n=8), Letrozole 0.3 mg/kg (n=8).

**Figure S2.**
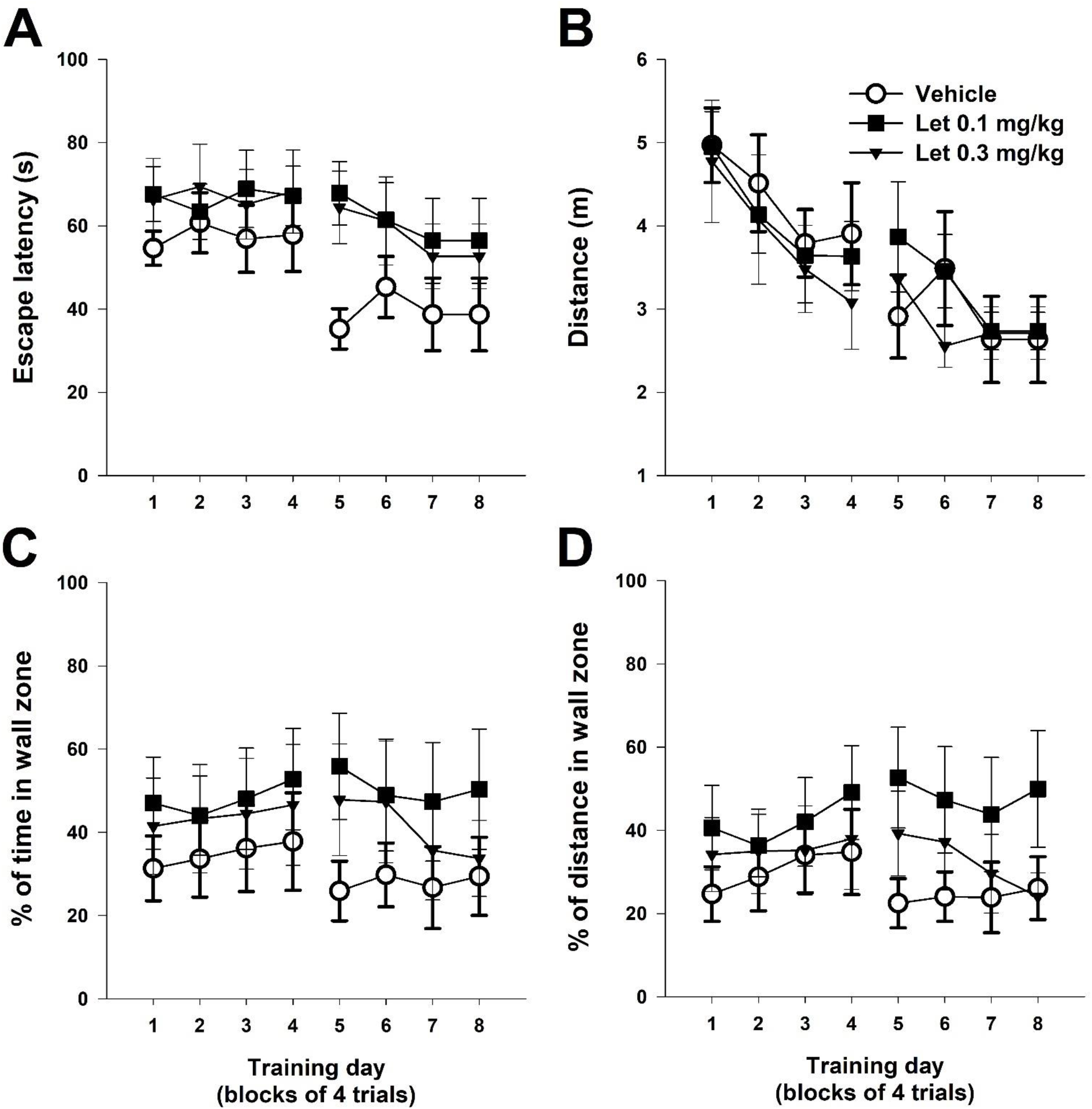
Learning performance of female BALB/c mice in the Morris water maze in a new room. Mice were previously trained in MWM in reference memory and reversal task for 4 weeks. Training was conducted during exposure to letrozole and following 8 weeks of prior exposure. Graphs show mean (A) escape latency and (B) mean distance mice swam during 4 trials in each training day to find a submerged platform; and (C) percentage of the escape latency and (D) percentage of the distance that a mouse spent/traveled in 10 cm wall zone in each trial. Data points and error bars represent mean and SEM, respectively. Vehicle (n=9), Letrozole 0.1 mg/kg (n=8), Letrozole 0.3 mg/kg (n=8).

**Figure S3.**
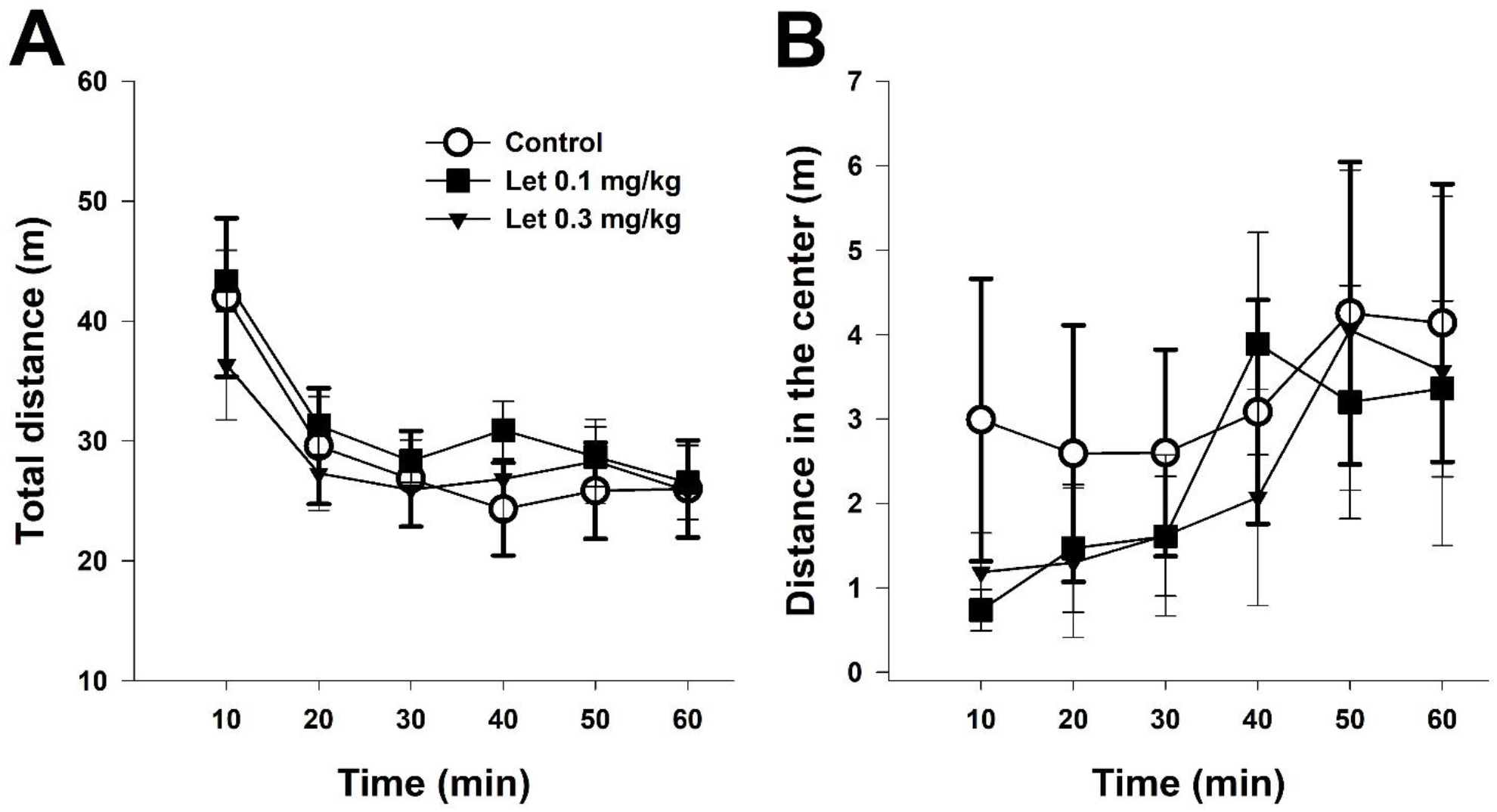
Exploration of the open field (120 cm × 120 cm) one week after completing the MWM training in a new room. Training was conducted during exposure to letrozole and following 11 weeks of prior exposure. Graphs show mean (A) total distance traveled and (B) distance traveled in the center of the arena (area outside the 20 cm wall zone). Data points and error bars represent mean and SEM, respectively. Vehicle (n=9), Letrozole 0.1 mg/kg (n=8), Letrozole 0.3 mg/kg (n=8).

**Figure S4.**
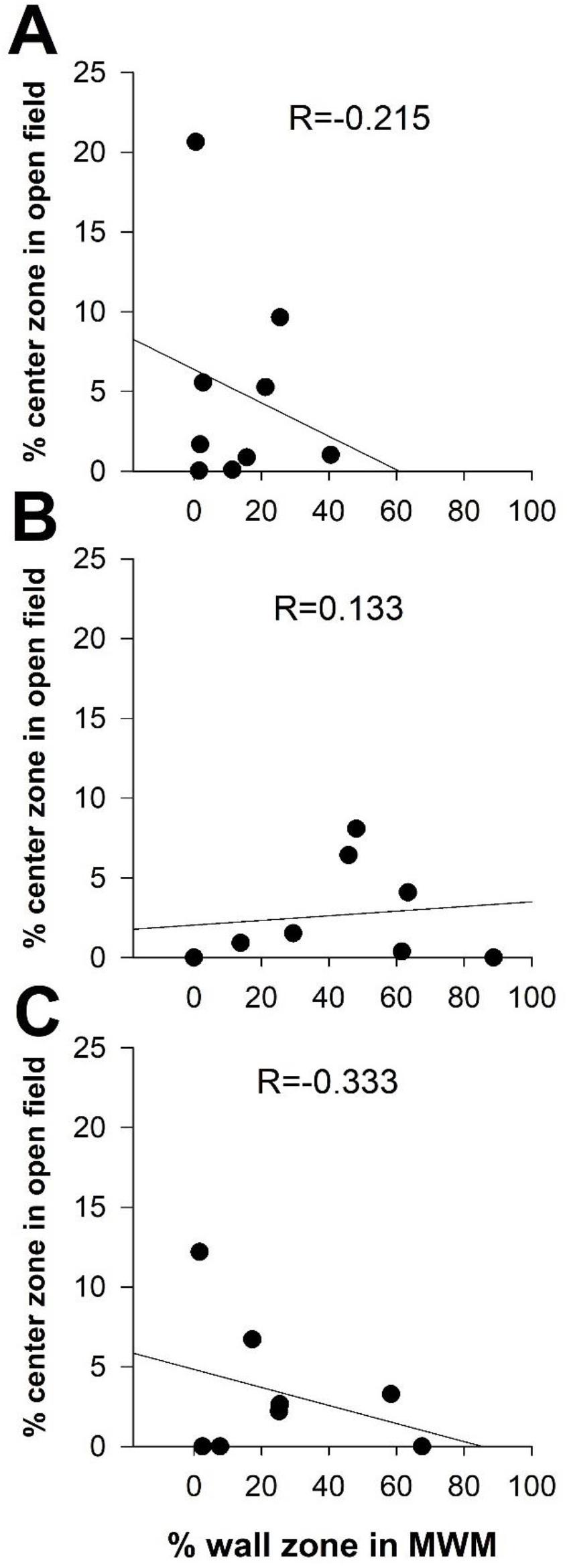
Correlation between mean % of distance in the MWM wall zone and mean % of distance traveled in the center of the open field arena (area outside the 20 cm wall zone). Vehicle (n=9), Letrozole 0.1 mg/kg (n=8), Letrozole 0.3 mg/kg (n=8).

